# The yeast proteases Ddi1 and Wss1 are both involved in the DNA replication stress response

**DOI:** 10.1101/584508

**Authors:** Michal Svoboda, Jan Konvalinka, Jean-François Trempe, Klara Grantz Saskova

**Author notes:** Corresponding author at: Institute of Organic Chemistry and Biochemistry of the Czech Academy of Sciences, Flemingovo n.2, 16610 Prague, Czech Republic, *E-mail address* (Klara Grantz Saskova).

## Abstract

Genome integrity and cell survival are dependent on proper replication stress response. Multiple repair pathways addressing obstacles generated by replication stress arose during evolution, and a detailed understanding of these processes is crucial for treatment of numerous human diseases. Here, we investigated the strong negative genetic interaction between two proteases involved in the DNA replication stress response, yeast Wss1 and Ddi1. While Wss1 proteolytically acts on DNA-protein crosslinks, mammalian DDI1 and DDI2 proteins remove RTF2 from stalled forks *via* a proposed proteasome shuttle hypothesis. We show that the double-deleted Δddi1, Δwss1 yeast strain is hypersensitive to the replication drug hydroxyurea and that this phenotype can be complemented only by catalytically competent Ddi1 protease. Furthermore, our data show the key involvement of the helical domain preceding the Ddi1 protease domain in response to replication stress caused by hydroxyurea, offering the first suggestion of this domain’s biological function. Overall, our study provides a basis for a novel dual protease-based mechanism enabling yeast cells to counteract DNA replication stress.

## 1. Introduction

During genome replication in eukaryotic cells, replication forks often encounter barriers that block progress down the DNA strand. These situations are generally referred to as replication stress (Zeman and Cimprich 2014), and multiple signaling pathways and processes responsible for counteracting replication stress arose during evolution (Branzei and Foiani 2009). As many replication-hindering barriers contain a protein component, it is unsurprising that proteases are involved in the cellular response to replication stress. The ubiquitin proteasome system plays a central role, with 26S proteasome involvement in regulated degradation of signaling molecules (Zhang et al. 2005), stalled RNA polymerases (Woudstra et al. 2002), and Ku70/80 proteins trapped after successful non-homologous end joining (van den Boom et al. 2016), as well as in general degradation of proteins trapped on the DNA strand (Larsen et al. 2018).

Researchers recently identified a novel proteolytic pathway acting in parallel with the ubiquitin proteasome system. This pathway involves direct proteolysis of proteins covalently trapped on the DNA strand to remove the replication fork-blocking barrier (Fielden et al. 2018; Stingele, Bellelli, and Boulton 2017). The process was described in *S. cerevisiae*, in which Wss1p, a DNA-activated metalloprotease, cleaves DNA-protein crosslinks (DPCs) such as compromised topoisomerases or products of formaldehyde toxicity in a replication-coupled manner (Balakirev et al. 2015; Stingele et al. 2014). After removal of the bulky protein, repair can continue through the nucleotide excision repair pathway. SPRTN, the mammalian orthologue of Wss1, acts as a DPC protease in mammalian cells, where it cleaves DPCs in a tightly coupled manner regulated by DNA-dependent and deubiquitination switches (Lopez-Mosqueda et al. 2016; Stingele et al. 2016; Vaz et al. 2016).

Other proteases recently found to be involved in the replication stress response include the Ddi1-like proteins. Mammalian DDI1 and DDI2 proteins associate with nascent replication forks and are involved in restarting stalled forks by facilitating removal of the replication termination factor RTF2 (Kottemann et al. 2018). DDI1- and DDI2-depleted cells fail to remove RTF2 from stalled forks, leading to accumulation of single-stranded DNA, sensitivity to replication drugs, and chromosome instability. Although both Ddi1-like proteins harbor a retroviral protease-like (RVP) domain, RTF2 removal has been proposed to proceed *via* a proteasomal shuttling mechanism. In proteasomal shuttling, proteins that contain ubiquitin-like (UBL) and ubiquitin-associated (UBA) domains serve as linkers between the 26S proteasome and polyubiquitinylated substrate (Finley 2009). This mechanism has been suggested for a yeast orthologue of Ddi1-like proteins, *S. cerevisiae* Ddi1p, which possesses UBL-UBA domain architecture (Fig. 1A) and serves as a proteasomal shuttle in degradation of HO endonuclease (Kaplun et al. 2005) and UFO1 protein (Ivantsiv et al. 2006; Voloshin et al. 2012). Yeast Ddi1p is inducible by DNA damage and shares a bidirectional promoter with MAG1, a DNA-3-methyladenine glycosylase involved in base-excision repair (Liu and Xiao 1997). Together with another proteasomal shuttle protein, Rad23, Ddi1p suppresses a temperature-sensitive mutation in Pds1 (securin), a mitotic checkpoint control protein, degradation of which by the UPS is required for mitosis (Clarke et al. 2001).

**Figure 1:**
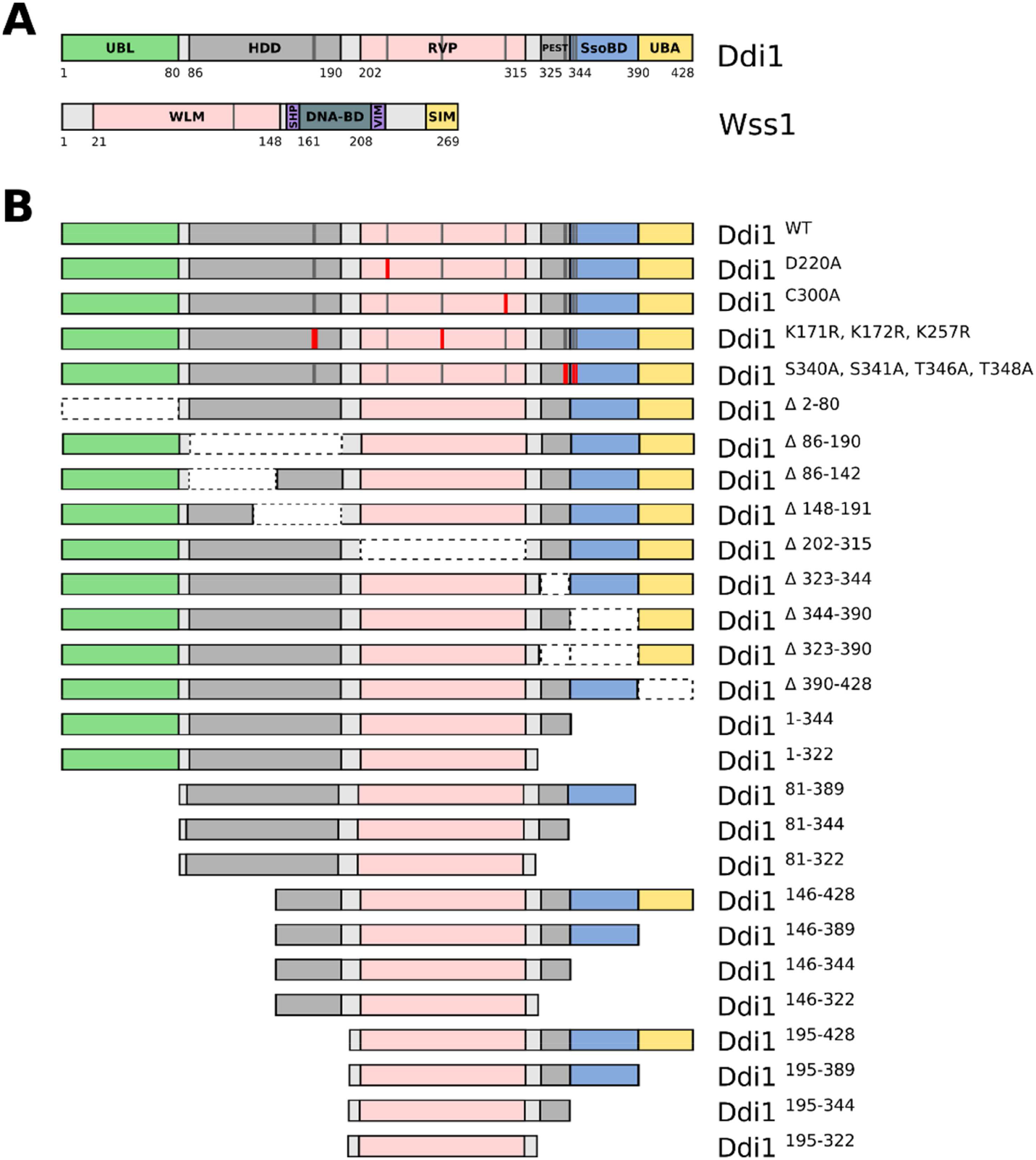
**A)** Schematic representation of the domain organization of *S. cerevisiae* Ddi1 and Wss1 proteins. UBL – ubiquitin-like domain, HDD – helical domain of Ddi1-like proteins, RVP – retroviral protease-like domain, PEST – proline, glutamate, serine and threonine rich domain targeting proteins for rapid degradation, SsoBD – Sso tSNARE binding domain, UBA – ubiquitin-associated domain, WLM – WSS1-like metalloprotease domain, SHP – Cdc48 interacting motif found in Shp1, DNA-BD – DNA-binding domain, VIM – VCP-interacting motif, SIM – SUMO-interacting motifs **B)** Point-mutated, domain-deleted and truncated constructs of *S. cerevisiae* Ddi1 used in this study. Truncated variants of Ddi1 protein were also prepared with C-terminal HA-tag.

A high-throughput synthetic lethality screen in yeast revealed a strong negative genetic interaction between *WSS1* and *DDI1* (Costanzo et al. 2016). The strong negative genetic interaction with another protease involved in the DNA replication stress response led us to question the proteasomal shuttling hypothesis for the role of Ddi1-like proteins in this response. In this study, we tested the sensitivity of a double-deleted yeast strain lacking *WSS1* and *DDI1* to DNA-damaging chemicals and identified a hypersensitivity to hydroxyurea, a ribonucleotide reductase inhibitor that induces nucleotide depletion and arrests cells in the S phase. Based on previous studies and our structural characterization of yeast and mammalian Ddi1/2 (Trempe et al. 2016; Siva et al. 2016), we performed rescue experiments with various deletions and single-site mutants of Ddi1 (Fig. 1B) to assess the involvement of particular yeast Ddi1 domains in the replication stress response.

## 2. Material and methods

### 2.1. Yeast strains and growth conditions

All *Saccharomyces cerevisiae* strains were isogenic derivatives of strain S288C, in the Y7092 background (*MATα can1Δ::STE2pr-Sp_his5 lyp1Δ ura3Δ0 leu2Δ0 his3Δ1 met15Δ0*), and were obtained as a kind gift from Prof. Charles Boone (Costanzo et al. 2016). Strain genotypes are listed in Supplementary Table S1. Standard yeast YPD (1% yeast extract, 2% bacto-peptone, 2% glucose) and SC or SC dropout media were used (Sherman 2002). For solid media, 1.5% agar was added. All cultivations were performed at 30 °C; liquid cultures were shaken at 260 RPM in an orbital shaker.

### 2.2. Plasmid construction and yeast transformation

Plasmids used for knockout phenotype rescue experiments were all derived from CEN-bound plasmid pAG416GPD-ccdB (a gift from Susan Lindquist, Addgene plasmid #14148) (Alberti, Gitler, and Lindquist 2007). The wild-type *Ddi1* sequence was amplified from the plasmid pET16_wt_Ddi1 (Trempe et al. 2016) using primers P1 and P2; the human *DDI2* sequence was amplified from pET16_wt_Ddi2 (Siva et al. 2016) using primers P45 and P46 (human DDI1 gene NM_001001711 was synthesized by GenScript and the coding sequence was amplified using primers P43 and P44). AttB adaptors were added by PCR with universal attB primers. Entry clones in pDONR221 were created using the standard protocol for Gateway BP clonase II (Thermo Fisher Scientific). Point mutants in the yeast *Ddi1* sequence were generated using the QuikChange Site-Directed Mutagenesis Kit (Agilent). Entry clones bearing domain deletions, truncated constructs and HA tagged constructs of yeast *Ddi1* were prepared according to the Gibson assembly procedure (Gibson 2011). Final expression plasmids for knockout phenotype rescue experiments were generated by recombination cloning from the given entry clone into pAG416GPD-ccdB, using Gateway LR clonase II (Thermo Fisher Scientific). All primers and plasmids are listed in Supplementary Table S2. All plasmids were verified by sequencing. Yeast were transformed using the standard LiAc/PEG3350 protocol (Gietz and Schiestl 2007).

### 2.3. Phenotypic characterization

Spot assays were performed by plating equal amounts of exponentially growing cells in serial dilutions onto solid YPDA media plates in the absence or presence of DNA-damage or DNA replication stress-causing substances. Plates were photographed after 60 hours of incubation at 30 °C.

### 2.4. Immunoblotting

Whole-cell protein extracts were prepared from exponentially growing cells as described (Kushnirov 2000). Proteins were separated in 12% Tris-Glycine SDS PAGE and wet-transferred onto a nitrocellulose membrane. Immunoblotting was performed using anti-PGK1 (1:5000, Novex 459250), anti-Ddi1 (1:5000, rabbit polyclonal, kind gift from Prof. Gerst, Weizmann Institute of Science (Lustgarten and Gerst 1999)) and C29F4 anti-HA (1:1000, Cell Signaling 3724) antibodies. Immunoblots were developed using near-infrared fluorophore-conjugated goat anti-mouse and anti-rabbit secondary antibodies (LI-COR 926-68070, 926-32211) and visualized with a LI-COR Odyssey CLx NIR fluorescence reader.

## 3. Results

### 3.1. Simultaneous loss of Ddi1 and Wss1 proteases leads to hypersensitivity to hydroxyurea

According to published data available in TheCellMap.org database (Usaj et al. 2017), the *DDI1* gene displays severe synthetic sickness when deleted together with the *WSS1* gene (Costanzo et al. 2016). As the protein products of both genes are known to be induced by DNA damage (Liu and Xiao 1997) or directly involved in the cellular response to DNA damage (Stingele et al. 2014), we decided to test the effects of various DNA-damaging chemicals on growth of a Δddi1, Δwss1 double-deleted yeast strain. Unlike the previously reported *DDI1* and *TDP1* double deletion, the *DDI1* and *WSS1* double deletion did not confer sensitivity to the topoisomerase inhibitor camptothecin (Fig.2). Hydrogen peroxide, which causes oxidative damage, also had no effect. The Δddi1, Δwss1 strain exhibited a slightly elevated sensitivity to methyl methane sulfonate and to long-term exposure to small concentrations of formaldehyde. However, the most pronounced phenotype of the double-deleted strain, compared to wild-type or single knock-out strains, was severe hypersensitivity to the ribonucleotide reductase inhibitor hydroxyurea (Fig. 2).

**Figure 2:**
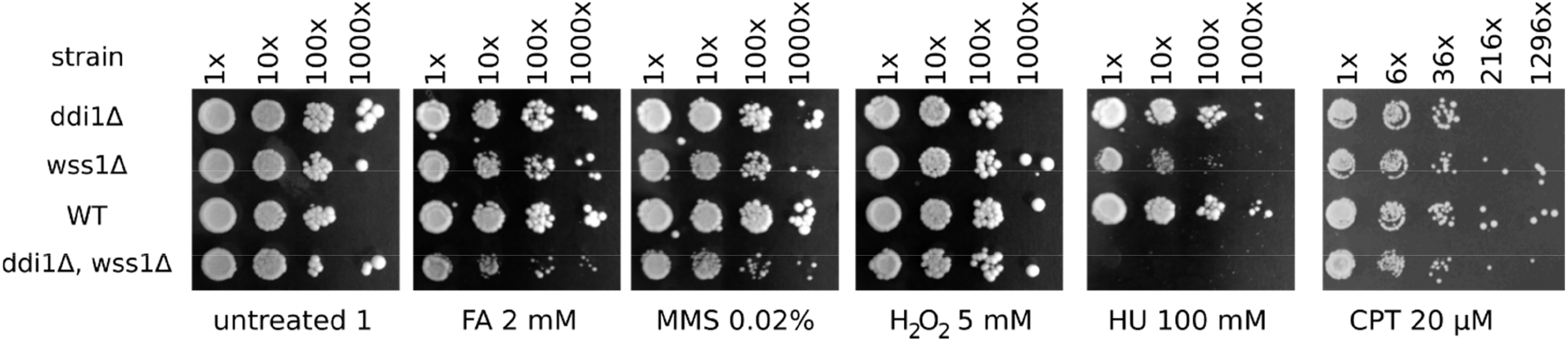
The Δddi1, Δwss1 strain is hypersensitive to hydroxyurea. Dilution spot assays testing the viability of wild-type, single knock-out and double knock-out strains under the influence of different DNA damage-causing chemicals. Equal amounts of exponentially growing cells were plated in serial dilutions on plates containing either YPDA media only or YPDA supplemented with formaldehyde (FA), methyl methane sulfonate (MMS), hydrogen peroxide, hydroxyurea (HU) or camptothecin (CPT).

### 3.2. Hydroxyurea hypersensitivity can be rescued by overexpression of wild-type Ddi1 but not a catalytically inactive mutant

The *DDI1* gene sequence contains activating elements for the *MAG1* gene, with which it shares a bidirectional promoter (Liu, Dai, and Xiao 1997). To test whether hydroxyurea hypersensitivity in the double-deletion strain was caused by the loss of Ddi1 expression rather than disruption of MAG1 expression, we performed a rescue experiment. The Δddi1, Δwss1 strain was transformed with a single copy of CEN-bound plasmid encoding wild-type Ddi1 protein expressed under the GPD constitutive promoter. Overexpression of wild-type protein led to complete rescue of the hydroxyurea hypersensitivity phenotype (Fig. 3A). This result confirms that the hypersensitivity to hydroxyurea observed in the Δddi1, Δwss1 strain is caused by the loss of the Ddi1 protein.

**Figure 3:**
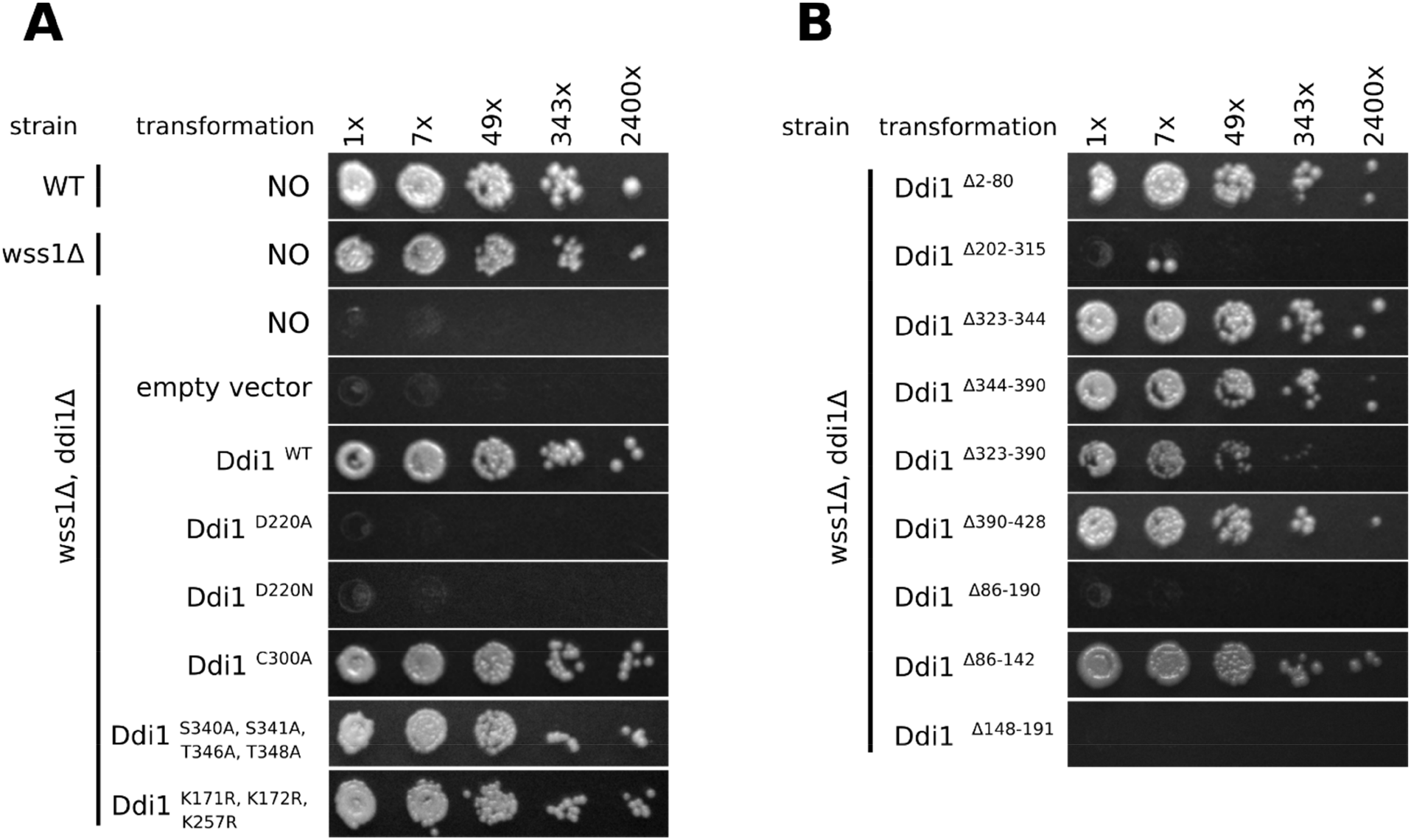
Retroviral protease-like (RVP) domain catalytic activity and the presence of a helical bundle (HDD) preceding the RVP domain are indispensable for complementation of hydroxyurea sensitivity. Equal amounts of exponentially growing cells transformed with single-copy plasmids overexpressing under control of the GPD promotor **A)** point-mutated variants of Ddi1 protein or **B)** variants with deletions of known structured domains of Ddi1 were plated in 7-fold serial dilutions on YPDA plates supplemented with 50 mM hydroxyurea.

Next, we explored the roles of several point mutations at Ddi1 residues of known functional importance. Mutation of all three lysine residues targeted by ubiquitination, as well as all four residues known to be phosphorylated during Ddi1’s involvement in exocytosis regulation (Gabriely et al. 2008), did not have an effect on the ability of the overexpressed protein to rescue hydroxyurea hypersensitivity, nor did mutation of Cys300, which was previously shown to influence Ddi1 nuclear localization (Gabriely et al. 2008). On the other hand, mutation of the catalytic aspartate (Asp220) in the RVP domain (Krylov and Koonin 2001; Sirkis, Gerst, and Fass 2006; Trempe et al. 2016) led to a complete loss of rescue, showing a phenotype identical to that of the original Δddi1, Δwss1 double mutant strain (Fig. 3A). Thus, resistance to hydroxyurea requires the enzymatic activity of the Ddi1 aspartyl protease domain.

### 3.3. The multi-helical bundle preceding the RVP domain is essential for Ddi1’s role in the DNA replication stress response

NMR and X-ray scattering experiments have shown that yeast Ddi1 is a multi-domain protein composed of multiple structured domains separated by flexible linkers (Trempe et al. 2016; Nowicka et al. 2015b). To test whether Ddi1 works in replication stress reversal as a whole protein or if only some of its domains are involved, we created plasmids encoding truncated variants of the protein lacking one particular domain (summarized in Fig. 1B). Expression of Ddi1 variants was confirmed by immunoblotting using either anti-Ddi1 antibody or anti-HA antibody (Fig. S1). Constructs lacking either the UBL (Δ2-80), PEST (Δ323-344), Sso binding (Δ344-390) or UBA (Δ390-428) domains were capable of fully rescuing Δddi1, Δwss1 hydroxyurea hypersensitivity (Fig. 3B). A construct lacking both the PEST and Sso binding domains (Δ323-390) performed less effectively. As expected from our finding for a catalytic Asp220 mutation, a Ddi1 variant lacking the RVP domain (Δ202-315) completely lost the ability to rescue hydroxyurea hypersensitivity. The same phenotype was also observed for Ddi1 lacking the recently identified Helical Domain of Ddi (HDD), a domain of unknown function (Siva et al. 2016; Trempe et al. 2016). Based on NMR structural data, HDD appears to comprise two multi-helical bundles connected by a flexible linker (Trempe et al. 2016). We therefore also prepared constructs lacking either the first or the second helical bundle of HDD. Only the second bundle spanning residues 148-191, located just before the protease domain, seems to be essential for Ddi1’s function in hydroxyurea hypersensitivity rescue.

### 3.4. Ubiquitin-like and ubiquitin-associated domains are not required for hydroxyurea hypersensitivity rescue

To further explore the mechanism of action of Ddi1 in the DNA replication stress response, we assessed which combination of domains is most essential for hydroxyurea hypersensitivity rescue. Neither the UBL nor UBA domain is required, as constructs lacking both (81-389) are capable of rescuing the phenotype, albeit less efficiently (Fig. 4). On the other hand, constructs with N-terminal truncation beyond the second helical bundle (195-xxx, etc.) were incapable of rescue, highlighting the crucial role of the second helical bundle in the DNA replication stress response. We identified a construct containing the second helical bundle and RVP domain (residues 146-322) as a minimal construct sufficient to fill Ddi1’s role in the DNA replication stress response, although we observed a slight decrease in effectivity. Due to loss of anti-Ddi1 antibody detection of some of the truncated variants, we performed identical dilution spot experiment using C-terminally HA-tagged Ddi1 protein variants and obtained the same results (Fig. S1).

**Figure 4:**
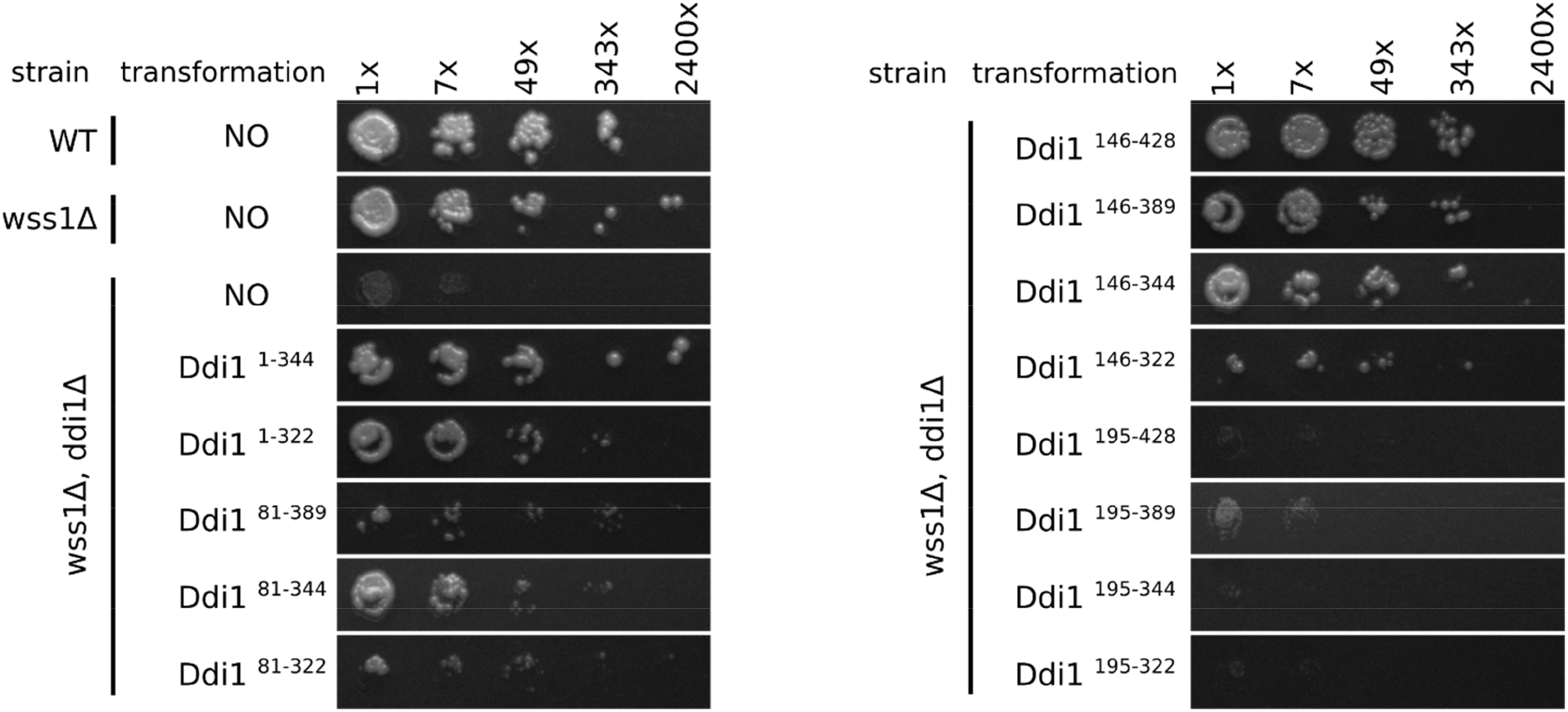
Complementation of hydroxyurea sensitivity by Ddi1 overexpression does not require the ubiquitin-like and ubiquitin-associated domains of Ddi1. The minimal construct capable of complementation spans residue 146 to residue 322, covering the multi-helical bundle and retroviral protease-like domain. Dilution spot assays with 7-fold serial dilutions of cells overexpressing truncated variants of Ddi1 protein plated on YPDA supplemented with 50 mM hydroxyurea are shown.

### 3.5. Human Ddi1-like proteins can partially rescue hydroxyurea hypersensitivity

Next, we tested whether our results also apply to human Ddi1 orthologues that contain the HDD helical domain preceding the retroviral aspartyl protease domain. We found that overexpression of both human DDI1 and DDI2 proteins can partially complement Δddi1, Δwss1 hydroxyurea sensitivity (Fig. 5), but the observed effect is weaker compared to yeast Ddi1 overexpression.

**Figure 5:**
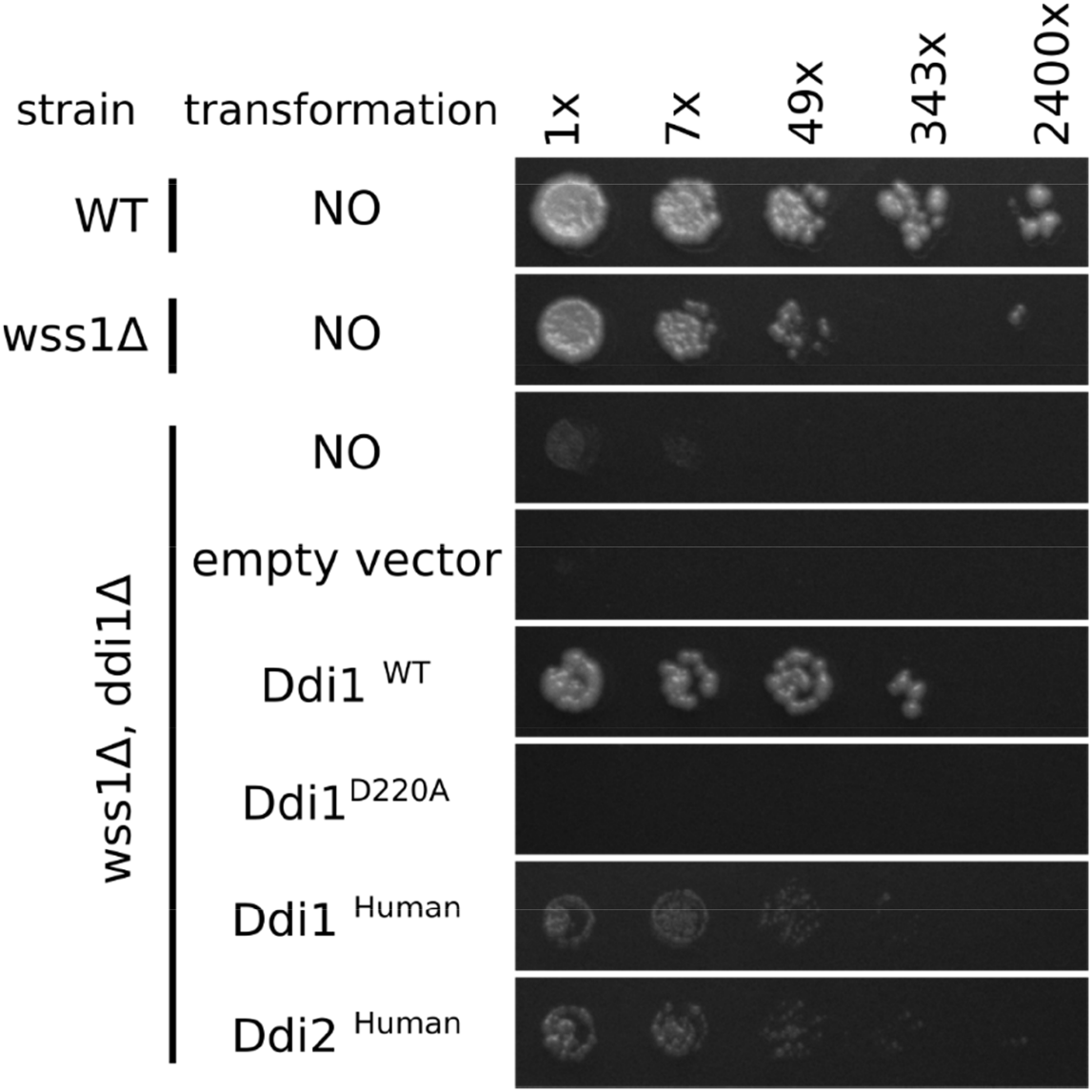
Human DDI1 and DDI2 proteins are capable of partial complementation of hydroxyurea sensitivity. Dilution spot assays with 7-fold serial dilutions of cells overexpressing yeast and human Ddi1-like proteins plated on YPDA supplemented with 50 mM hydroxyurea are shown.

## 4. Discussion

The ability to counter DNA replication stress by repairing and restarting stalled replication forks is essential for successful completion of the cell cycle. Therefore, multiple pathways have evolved to respond to replication stress, including the activity of proteolytic enzymes. This study focuses on the characterization of the role of yeast Ddi1 protein in the DNA replication stress response. As part of a genome-wide synthetic lethality screen in yeast, Ddi1 exhibited a strong negative genetic interaction with Wss1, another replication stress-countering protease (Costanzo et al. 2016). Another study identified human proteins DDI1 and DDI2, mammalian orthologues of Ddi1 that are involved in degradation of the replication termination factor RTF2, facilitating restart of stalled replication forks (Kottemann et al. 2018).

Inspired by the Synthetic Genetic Array in yeast, we decided to explore the interplay between Ddi1 and Wss1 with a focus on their role in response to DNA damage and DNA replication stress. By testing the sensitivity of the double-mutated Δddi1, Δwss1 yeast strain to various DNA damage agents, we found the strain to be extremely sensitive to hydroxyurea. Hydroxyurea causes replication fork stalling by inhibiting ribonucleotide reductase and therefore depleting free nucleotide triphosphate building blocks. Unlike in mammalian cells, Ddi1 deletion in yeast does not itself cause sensitivity to hydroxyurea. Deletion of Wss1 had been previously found to cause mild sensitivity to hydroxyurea (O’Neill et al. 2004), as confirmed in our experiments. The synthetic effect of both mutations substantially affected hydroxyurea sensitivity, beyond simple additivity. This suggests that Ddi1 and Wss1 act in two independent pathways, each counteracting the replication stress caused by hydroxyurea. As hydroxyurea is not known to directly induce DPCs, the main type of lesion resolved by Wss1, our observation raises a number of questions. For example, it remains unclear which DNA damage response pathway triggered by hydroxyurea leads to growth arrest when both Ddi1 and Wss1 are lost. The double deletion strain may promote upregulation of checkpoint pathways in the S phase, which induce cell cycle arrest in the presence of DNA damage. Other questions to be addressed included which substrate(s) the Ddi1 protease domain cleaves in this context and whether the target molecule is the same for Wss1.

Next, we focused on understanding the mechanism of Ddi1’s function in the yeast replication stress response. First, we confirmed that the observed hydroxyurea sensitivity is indeed caused by the loss of Ddi1 expression by overexpressing Ddi1 from a plasmid to complement the phenotype. Next, we analyzed Ddi1’s mode of action by complementation experiments with several Ddi1 variants derived from previous functional and structural studies. We found that the catalytic activity of the RVP domain is essential. Indeed, there were only two specific complementation variants that completely abolished rescue of the hydroxyurea sensitivity phenotype: mutation in the catalytic aspartate of RVP and deletion of the helical bundle directly preceding RVP. Our data are thus consistent with a previous observation (Gabriely et al. 2008) identifying the catalytically active Ddi1 protease domain as necessary for the rescue of *pds1-128* cells. Many publications describing yeast Ddi1 have focused on its role as a proteasomal shuttling protein, bringing polyubiquitinated substrates to the 26S proteasome (Clarke et al. 2001; Finley 2009; Ivantsiv et al. 2006; Kaplun et al. 2005; Voloshin et al. 2012). A shuttling mechanism has also been proposed for human Ddi1-like proteins in the degradation of RTF2 and subsequent restart of replication forks (Kottemann et al. 2018). In contrast, our data do not support the proteasome shuttling mechanism, as deletion of UBL, UBA or both did not impair the ability of the protein to rescue hydroxyurea sensitivity. Our findings also contrast with the so-called alternative proteasomal shuttle hypothesis proposed by Nowicka and coworkers (Nowicka et al. 2015a), which suggests that the first Ddi1 UBL domain binds polyubiquitinated substrate and the other UBL targets the proteasome. The hydroxyurea hypersensitivity phenotype of the double-mutated Δddi1, Δwss1 yeast strain can be partially complemented even with a minimal construct comprising the double helical bundle followed by the RVP domain (Ddi1 146-322).

To assess whether our results also apply to human orthologues, we complemented the double-deleted yeast strain with plasmids encoding human DDI1 and DDI2 wild-type proteins. Both human DDI1 orthologues showed partial complementation of hydroxyurea sensitivity. This can be explained by differences in the structure of putative substrates (RTF2, a known target of both human Ddi1-like proteins, does not have an orthologue in *S. cerevisiae*) as well as by reasons rooted in enzymology. Proteases usually have a relatively narrow temperature optimum, and the 7 °C temperature difference between the temperature optima of most human enzymes and cultivation temperature for yeast cultures may be sufficient to impair the activity of the human Ddi1 proteases. Nevertheless, the experiment demonstrates a conservation of function between yeast and human Ddi1 orthologues. Notably, sequence conservation in Ddi1 orthologues is higher for the second bundle of the HDD and the protease domains, and some orthologues lack either the UBL or the UBA domain (Siva et al. 2016; Trempe et al. 2016). This is consistent with the main function of Ddi1 being dependent on its protease activity, perhaps with the substrate-recognition function carried out by the HDD domain.

Taken together, our data suggest the existence of a dual protease mechanism providing yeast cells with the ability to overcome DNA-replication stress caused by the ribonucleotide reductase inhibitor hydroxyurea. Furthermore, we identified a putative function for the recently described HDD domain. The substrate repertoire of these proteases and the detailed mechanistic basis of this new pathway remain to be further analyzed.

## Acknowledgements

We thank Drs. Jason Tanny, Jan Šilhán and Monika Sivá for useful discussions. We thank Prof. Charles Boone for providing yeast strains, Prof. Jeffrey E. Gerst for anti-Ddi1 antibody and Prof. Susan Lindquist for pAG416GPD-ccdB plasmid. We also thank Dr. Hillary Hoffman for language editing.

