## Supplementary Material for "The yeast proteases Ddi1 and Wss1 are both involved in the DNA replication stress response"

**Supplementary Table S1:** *S.cerevisiae* strains used in this study

| Name | Strain | Relevant Genotype (S288c) | Source |
|---|---|---|---|
| background | Y7092 | *MATα, can1Δ0::STEpr-Sp_His5, lyp1Δ0, his3Δ1, leu2Δ0, ura3Δ0, met15Δ0* | (Costanzo et al. 2016) |
| Δddi1 | YER143W_sn475 | *Y7092, ddi1Δ::natMX4* | (Costanzo et al. 2016) |
| Δwss1 | YHR134W_sn1569 | *Y7092, wss1Δ::natMX4* | (Costanzo et al. 2016) |
| WT | Y8835 | *Y7092, ura3Δ0::natMX4* | (Costanzo et al. 2016) |
| Δddi1, Δwss1 | | *Y7092, MATa, ddi1Δ::natMX4, wss1Δ::kanMX4* | (Costanzo et al. 2016) |

**Supplementary Table S2:** Plasmids and primers used in this study

| Name | Code | Relevant Sequence | Source; Primers used |
|---|---|---|---|
| empty vector | pAG14148 | pAG416GPD-ccdB | (Alberti, Gitler, and Lindquist 2007) |

| | | | |
|---|---|---|---|
| pENTRY | vMSV001 | pDONR221- Ddi1$^{WT}$ | this study; P1, P2 |
| Ddi1$^{WT}$ | vMSV076 | pAG416GPD-Ddi1$^{WT}$ | this study; P1, P2 |
| Ddi1$^{D220A}$ | vMSV077 | pAG416GPD-Ddi1$^{D220A}$ | this study; P3, P4 |
| Ddi1$^{\Delta 2\text{-}80}$ | vMSV078 | pAG416GPD-Ddi1$^{\Delta 2\text{-}80}$ | this study; P5, P6 |
| Ddi1$^{\Delta 390\text{-}428}$ | vMSV079 | pAG416GPD-Ddi1$^{\Delta 390\text{-}428}$ | this study; P7, P8 |
| Ddi1$^{\Delta 344\text{-}390}$ | vMSV080 | pAG416GPD-Ddi1$^{\Delta 344\text{-}390}$ | this study; P9, P10 |
| Ddi1$^{\Delta 323\text{-}344}$ | vMSV081 | pAG416GPD-Ddi1$^{\Delta 323\text{-}344}$ | this study; P11, P12 |
| Ddi1$^{\Delta 323\text{-}390}$ | vMSV082 | pAG416GPD-Ddi1$^{\Delta 323\text{-}390}$ | this study; P13, P14 |
| Ddi1$^{\Delta 202\text{-}315}$ | vMSV083 | pAG416GPD-Ddi1$^{\Delta 202\text{-}315}$ | this study; P15, P16 |
| Ddi1$^{\Delta 86\text{-}190}$ | vMSV084 | pAG416GPD-Ddi1$^{\Delta 86\text{-}190}$ | this study; P17, P18 |
| Ddi1$^{81\text{-}389}$ | vMSV085 | pAG416GPD-Ddi1$^{81\text{-}389}$ | this study; P5, P6, P7, P8 |
| Ddi1$^{K171R,\ K172R,\ K257R}$ | vMSV092 | pAG416GPD-Ddi1$^{K171R,\ K172R,\ K257R}$ | this study; P19, P20, P21, P22, P23, P24 |
| Ddi1$^{S340A,\ S341A,\ T346A,\ T348A}$ | vMSV095 | pAG416GPD-Ddi1$^{S340A,\ S341A,\ T346A,\ T348A}$ | this study; P25, P26, P27, P28, P29, P30, P31, P32 |
| Ddi1$^{\Delta 86\text{-}142}$ | vMSV096 | pAG416GPD-Ddi1$^{\Delta 86\text{-}142}$ | this study; P33, P34 |
| Ddi1$^{\Delta 148\text{-}191}$ | vMSV097 | pAG416GPD-Ddi1$^{\Delta 148\text{-}191}$ | this study; P35, P36 |
| Ddi1$^{195\text{-}322}$ | vMSV098 | pAG416GPD-Ddi1$^{195\text{-}322}$ | this study; P6, P7, P37, P38 |
| Ddi1$^{146\text{-}428}$ | vMSV099 | pAG416GPD-Ddi1$^{146\text{-}428}$ | this study; P6, P40 |
| Ddi1$^{195\text{-}428}$ | vMSV100 | pAG416GPD-Ddi1$^{195\text{-}428}$ | this study; P6, P37 |
| Ddi1$^{1\text{-}344}$ | vMSV101 | pAG416GPD-Ddi1$^{1\text{-}344}$ | this study; P7, P39 |
| Ddi1$^{1\text{-}322}$ | vMSV102 | pAG416GPD-Ddi1$^{1\text{-}322}$ | this study; P7, P38 |
| Ddi1$^{146\text{-}389}$ | vMSV103 | pAG416GPD-Ddi1$^{146\text{-}389}$ | this study; P6, P7, P8, P40 |
| Ddi1$^{195\text{-}389}$ | vMSV104 | pAG416GPD-Ddi1$^{195\text{-}389}$ | this study; P6, P7, P8, P37 |
| Ddi1$^{81\text{-}344}$ | vMSV105 | pAG416GPD-Ddi1$^{81\text{-}344}$ | this study; P5, P6, P7, P39 |
| Ddi1$^{146\text{-}344}$ | vMSV106 | pAG416GPD-Ddi1$^{146\text{-}344}$ | this study; P6, P7, P39, P40 |

| | | | |
|---|---|---|---|
| Ddi1$^{195-344}$ | vMSV107 | pAG416GPD-Ddi1$^{195-344}$ | this study; P6, P7, P37, P39 |
| Ddi1$^{81-322}$ | vMSV108 | pAG416GPD-Ddi1$^{81-322}$ | this study; P5, P6, P7, P38 |
| Ddi1$^{146-322}$ | vMSV109 | pAG416GPD-Ddi1$^{146-322}$ | this study; P6, P7, P38, P40 |
| Ddi1$^{C300A}$ | vMSV114 | pAG416GPD-Ddi1$^{C300A}$ | this study; P41, P42 |
| Ddi1$^{human}$ | vMSV115 | pAG416GPD-Ddi1$^{human}$ | this study; P43, P44 |
| Ddi2$^{human}$ | vMSV116 | pAG416GPD-Ddi2$^{human}$ | this study; P45, P46 |
| Ddi1_HA$^{81-389}$ | vMSV125 | pAG416GPD-Ddi1_HA$^{81-389}$ | this study; P5, P6, P47, P48 |
| Ddi1_HA$^{81-344}$ | vMSV126 | pAG416GPD-Ddi1_HA$^{81-344}$ | this study; P5, P6, P47, P49 |
| Ddi1_HA$^{146-389}$ | vMSV127 | pAG416GPD-Ddi1_HA$^{146-389}$ | this study; P6, P40, P47, P48 |
| Ddi1_HA$^{146-344}$ | vMSV128 | pAG416GPD-Ddi1_HA$^{146-344}$ | this study; P6, P40, P47, P49 |
| Ddi1_HA$^{146-322}$ | vMSV129 | pAG416GPD-Ddi1_HA$^{146-322}$ | this study; P6, P40, P47, P50 |
| Ddi1_HA$^{195-389}$ | vMSV130 | pAG416GPD-Ddi1_HA$^{195-389}$ | this study; P6, P37, P47, P48 |
| Ddi1_HA$^{195-344}$ | vMSV131 | pAG416GPD-Ddi1_HA$^{195-344}$ | this study; P6, P37, P47, P49 |
| Ddi1_HA$^{195-322}$ | vMSV132 | pAG416GPD-Ddi1_HA$^{195-322}$ | this study; P6, P37, P47, P50 |

| Primer | Sequence (5´-3´) |
|---|---|
| P1 | AAAAAGCAGGCTACAAAATGGATTTAACAATTTCAAACG |
| P2 | AGAAAGCTGGGTTCTATCATTGGAAAAGGAGGGATG |
| attB1 uni | GGGGACAAGTTTGTACAAAAAAGCAGGCT |
| attB2 uni | GGGGACCACTTTGTACAAGAAAGCTGGGT |
| P3 | AAAGGCATTTGTAGCTACAGGGGCTCAAA |
| P4 | TTTGAGCCCCTGTAGCTACAAATGCCTTT |
| P5 | GCAGGCTACAAAATGATTCAAACAGATGCTGCTACTTTG |
| P6 | CATTTTGTAGCCTGCTTTTTTGTAC |
| P7 | TGATAGAACCCAGCTTTCTTGTAC |
| P8 | AGCTGGGTTCTATCACGTTCTCCCGGTTGCCGTTG |

| | |
|---|---|
| P9 | GTCTGATAAGCCCGAACAAACGATTAAACAG |
| P10 | TCGGGCTTATCAGACGAAGTTGTAACTGAC |
| P11 | GAAGCGGAACTAACACCCACCAAGACTAG |
| P12 | GTGTTAGTTCCGCTTCACTCAAAAAGC |
| P13 | GGAACCCGAACAAACGATTAAACAG |
| P14 | GTTTGTTCGGGTTCCGCTTCACTCAAAAAGC |
| P15 | ACCCAGGTCAGCTTTTTGAGTGAAGCGG |
| P16 | AAAGCTGACCTGGGTAAACATTTCAGG |
| P17 | ACAGATGCTATCGAATATACACCTGAAATGTTTACC |
| P18 | TATTCGATAGCATCTGTTTGAATGGAATTG |
| P19 | ATGATCCTGACAACAGGAAGAGGATTGCAGA |
| P20 | TCTGCAATCCTCTTCCTGTTGTCAGGATCAT |
| P21 | ATCCTGACAACAGGAGGAGGATTGCAGAGCT |
| P22 | AGCTCTGCAATCCTCCTCCTGTTGTCAGGAT |
| P23 | GCGTAGGAACCGGCAGAATTATTGGGAGAAT |
| P24 | ATTCTCCCAATAATTCTGCCGGTTCCTACGC |
| P25 | TCAGTTACAACTTCGGCTGATAAGCCCCTA |
| P26 | TAGGGGCTTATCAGCCGAAGTTGTAACTGA |
| P27 | TCAGTTACAACTGCGGCTGATAAGCCCCTA |
| P28 | TAGGGGCTTATCAGCCGCAGTTGTAACTGA |
| P29 | CTGATAAGCCCCTAGCACCCACCAAGACTAG |
| P30 | CTAGTCTTGGTGGGTGCTAGGGGCTTATCAG |
| P31 | CTGATAAGCCCCTAGCACCCGCCAAGACTAG |
| P32 | CTAGTCTTGGCGGGTGCTAGGGGCTTATCAG |
| P33 | ACAGATGCTGGCTATAACACCGCCATG |
| P34 | GTTATAGCCAGCATCTGTTTGAATGGAATTG |
| P35 | CACCGCCGAATATACACCTGAAATGTTTACCCAGGTC |
| P36 | AGGTGTATATTCGGCGGTGTTATAGCCACCATAAC |

| | |
|---|---|
| P37 | GCAGGCTACAAAATGCCTGAAATGTTTACCCAGGTC |
| P38 | AGCTGGGTTCTATCATTCCGCTTCACTCAAAAAGC |
| P39 | AGCTGGGTTCTATCAGGGCTTATCAGACGAAGTTG |
| P40 | GCAGGCTACAAAATGACCGCCATGAATCCTTTTG |
| P41 | GCATTTGGCTGCTGTGGACTTAAAGGAAAAC |
| P42 | GTTTTCCTTTAAGTCCACAGCAGCCAAATGC |
| P43 | GCAGGCTACAAAATGCTGATCACCGTGTACTG |
| P44 | AGCTGGGTTCTATCAATGCTCTTTTCGTCCTGAATC |
| P45 | GCAGGCTACAAAATGCTGCTCACCGTGTACTG |
| P46 | AGCTGGGTTCTATCATGGCTTCTGACGCTCTGC |
| P47 | GGATCCTATCCATATGACGTTCCAGATTACGCTTGAT AGAACCCAGCTTTCTTG |
| P48 | ATATGGATAGGATCCCGTTCTCCCGGTTGCCGTTG |
| P59 | ATATGGATAGGATCCGGGCTTATCAGACGAAGTTG |
| P50 | ATATGGATAGGATCCTTCCGCTTCACTCAAAAAGC |

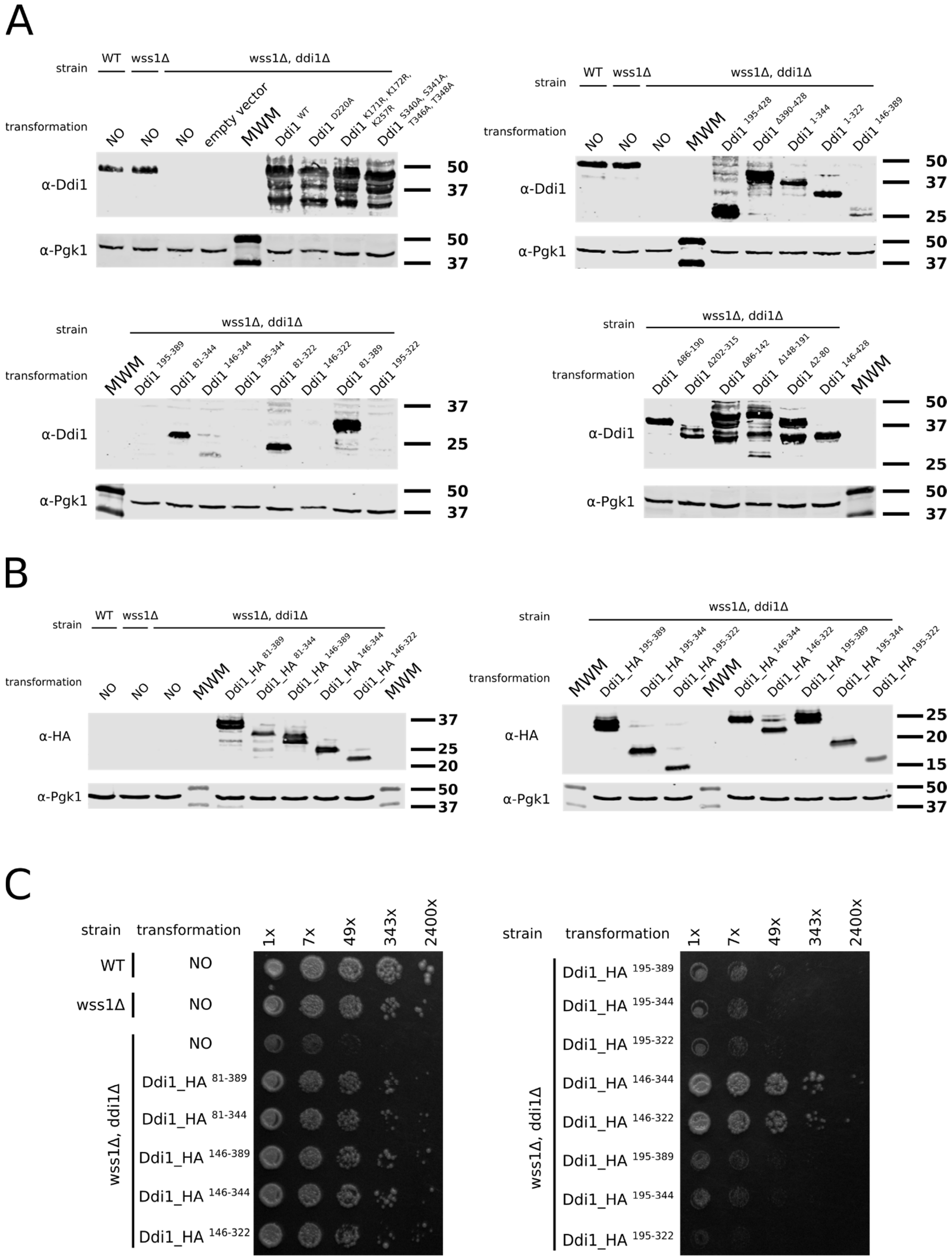
A
strain
WT
wss1Δ
wss1Δ, ddi1Δ
transformation
NO
empty vector
MWM
Ddi1 WT
Ddi1 D220A
Ddi1 K171R, K172R, K257R
Ddi1 S340A, S341A, T346A, T348A
α-Ddi1
α-Pgk1
50
37
Ddi1 195-428
Ddi1 Δ390-428
Ddi1 1-344
Ddi1 1-322
Ddi1 146-389
25
Ddi1 195-389
Ddi1 81-344
Ddi1 146-344
Ddi1 195-344
Ddi1 81-322
Ddi1 146-322
Ddi1 81-389
Ddi1 195-322
Ddi1 Δ86-190
Ddi1 Δ202-315
Ddi1 Δ86-142
Ddi1 Δ148-191
Ddi1 Δ2-80
Ddi1 146-428
B
Ddi1_HA 81-389
Ddi1_HA 81-344
Ddi1_HA 146-389
Ddi1_HA 146-344
Ddi1_HA 146-322
Ddi1_HA 195-389
Ddi1_HA 195-344
Ddi1_HA 195-322
α-HA
20
15
C
1x
7x
49x
343x
2400x
wss1Δ, ddi1Δ

**Supplementary Figure S1: A)** Verification of overexpression of Ddi1 protein variants used in this study. The whole yeast cell extracts were separated by 12% SDS-PAGE followed by Western blotting. Ddi1 detection was performed using anti-Ddi1 antibody (1:5000, rabbit polyclonal, kind gift from Prof. Gerst, Weizmann Institute of Science (Lustgarten and Gerst 1999)); Pgk1 (anti-Pgk1, 1:5000, Novex 459250) was used as a loading control. Size markers (in kDa) are indicated on the right side of the blots. In some cases, overexpressed Ddi1 clearly undergoes partial digestion. We were repeatedly unable to detect some of the Ddi1 variants (Ddi1 $^{146\text{-}322}$, Ddi1 $^{195\text{-}344}$, Ddi1 $^{195\text{-}389}$ and Ddi1 $^{195\text{-}322}$), most likely due to the loss of the Ddi1 antibody sensitivity because of the protein truncations. **B)** Truncated Ddi1 protein variants were prepared with C-terminal HA-tag and their overexpression was confirmed by Western blotting. Ddi1 detection was performed using rabbit monoclonal C29F4 anti-HA antibody (1:1000, Cell Signaling 3724); Pgk1 (anti-Pgk1, 1:5000, Novex 459250) was used as a loading control. Size markers (in kDa) are indicated on the right side of the blots. **C)** Dilution spot assays with 7-fold serial dilutions of cells overexpressing HA-tagged truncated variants of Ddi1 protein plated on YPDA supplemented with 50 mM hydroxyurea are shown. Results are in agreement with complementation study using truncated variants of Ddi1 protein lacking the C-terminal HA-tag.
